# Dysregulation of synaptic transcripts underlies network abnormalities in ALS patient-derived motor neurons

**DOI:** 10.1101/2024.05.29.596436

**Authors:** Anna M. Kollstrøm, Nicholas Christiansen, Axel Sandvig, Ioanna Sandvig

## Abstract

Amyotrophic lateral sclerosis (ALS) is characterized by dysfunction and loss of upper and lower motor neurons. Several studies have identified structural and functional alterations in the motor neurons before the manifestation of symptoms, yet the underlying cause of such alterations and how they contribute to the progressive degeneration of affected motor neuron networks remain unclear. Importantly, the short and long-term spatiotemporal dynamics of neuronal network activity make it challenging to discern how ALS-related network reconfigurations emerge and evolve. To address this, we systematically monitored the structural and functional dynamics of motor neuron networks with a confirmed endogenous C9orf72 mutation. We show that ALS patient-derived motor neurons display time-dependent neural network dysfunction, specifically reduced firing rate and spike amplitude, impaired bursting, but higher overall synchrony in network activity. These changes coincided with altered neurite outgrowth and branching within the networks. Moreover, transcriptional analyses revealed dysregulation of molecular pathways involved in synaptic development and maintenance, neurite outgrowth and cell adhesion, suggesting impaired synaptic stabilization. This study identifies early synaptic dysfunction as a contributing mechanism resulting in network-wide structural and functional compensation, which may over time render the networks vulnerable to neurodegeneration.

## Introduction

ALS is a complex disease involving a multitude of disease mechanisms ultimately resulting in retraction of the motor neuron axon from the neuromuscular junction, and cell death. Despite intensive efforts over the past decades to unravel the genetic and pathophysiological underpinnings of the disease, a complete understanding of disease etiology is still lacking. Consequently, effective treatment options for patients are lacking, and the disease results in rapidly progressing paralysis and death within 3-5 years after diagnosis.

Structurally, ALS primarily affects the upper motor neurons in the primary motor cortex and lower motor neurons in the brain-stem and spinal cord, but there is evidence of involvement of extra motor regions as well, including atrophy of frontotemporal areas (1) and progression of pTDP-43 pathology into prefrontal regions and the thalamus (2, 3). Importantly, these structural alterations are accompanied by changes in functional connectivity and neural activity (4, 5, 6). However, the specific relationship between structural impairment and disruptions in network communication remains elusive, particularly how such disruptions manifest at early stages of the disease, and to what extent they contribute to disease progression.

While some studies have demonstrated that increased functional connectivity in motor, prefrontal and thalamic regions correlate with disease progression (7, 8), increased connectivity in motor areas has also been identified in pre-symptomatic carriers of ALS-causing mutations, suggesting that functional changes is an early feature of the disease and not merely a consequence of atrophy (9, 10). Functional alterations in ALS have also been shown to involve cortical hyperexcitability (11). Notably, this has also been identified in pre-symptomatic individuals, suggesting a role in early stages of ALS pathogenesis (12, 13). However, contradictory findings point to hypoexcitability as a more likely mechanism contributing to motor neuron degeneration (14, 15). Accordingly, the exact nature of functional alterations in ALS, and the consequences thereof, are not clear.

This may be explained in part by the heterogeneity of the disease in terms of genetic mutations, family history, site of onset and disease duration (16). However, spatiotemporal aspects of functional network dynamics further complicate the identification of subtle network responses to progressive pathology. Crucially, as such early responses do not manifest as symptoms or direct impairment, investigations of the micro- and mesoscale level cellular and network responses are particularly challenging in patients and animal models. Thus, a comprehensive investigation of structural alterations and the corresponding early-onset functional responses in ALS must also consider the temporal dynamics of these relationships.

In this context, the self-organizing properties of neurons can be exploited to engineer complex neuronal networks which preserve hallmarks of network function in the brain, including emerging spontaneous activity of increasing spatiotemporal complexity (17, 18, 19). We and others have demonstrated that the application of such advanced cellular models enable the investigation of neural network dynamics associated with ALS pathology (20, 21, 22, 23, 24, 25, 26). Nevertheless, the role of such dynamics in ALS pathogenesis, especially those pertaining to neuronal excitability, remains elusive. Wainger et al. (21) observed increased spontaneous firing in SOD1-, C9orf72- and FUS-mutant hiPSC-derived motor neurons, whereas reduced excitability was found by both Sareen et al. (20) and Naujock et al. (22). Time-dependent shifts from a hyper- to a hypoexcitable state have also been reported (23, 24, 25), thereby emphasizing the importance of evolving network dynamics over time in response to pathology. Nevertheless, previous studies are characterized by temporally limited and infrequent investigations, thereby failing to capture the full complexity of neural network dynamics (27).

Here we aimed to identify how ALS-related disturbances to network activity emerge and evolve, and the molecular underpinnings contributing to such disturbances. We longitudinally investigated the structural, functional and transcriptional maturation of hiPSC-derived motor neuron networks with an endogenous *C9orf72* mutation. Both healthy control and ALS networks developed hallmarks of structural and functional network maturity. Nevertheless, motor neuron networks harboring the genetic mutation exhibited time-dependent disturbances in network activity, including reduced firing rate and spike amplitude, impaired bursting and increased overall synchrony in network activity. These functional disturbances were accompanied by neurite outgrowth abnormalities. Furthermore, transcriptional analyses revealed consistent dysregulation of synaptic and cell adhesion pathways. Collectively, our findings demonstrated that motor neurons with an endogenous ALS mutation harbored intrinsic disturbances to synaptic stabilization, which over time resulted in structural and functional alterations.

## Results

### Healthy and ALS patient-derived motor neuron networks display time-dependent differences in spontaneous activity

Longitudinal MEA recordings from 24 to 52 DIV revealed that both ALS patient-derived and healthy motor neuron networks exhibited a steady increase in spontaneous activity over time. Despite some network-to-network variation, both groups followed a similar development in the firing rate until 38 DIV, and no significant differences between the groups were identified (Fig. 1A). At 40 DIV, the ALS motor neuron networks displayed a significantly lower firing rate compared to that of healthy networks (p<.001). The firing rate remained lower in the ALS networks until the final recording at 52 DIV, and the increase in firing rate over time was not as prominent as for the healthy networks (see table S1 for p-values at all recording time points). The mean spike amplitude was significantly lower for the ALS networks at most recording time points, with the exception of four recordings from 32-40 DIV (Fig. 1B, table S1). Interestingly, from 40 DIV onward, the mean spike amplitude appeared to stabilize in the ALS networks, while it continued to increase over time for the healthy networks. The mean ISI decreased over time in both the ALS and the healthy networks, in correspondence with an increased firing rate, and developed in a similar manner between the two groups, although significantly higher in the ALS networks from 40 DIV onward (Fig. 1C; table S1). The coherence index was significantly higher in the ALS networks across all time points except for 28 DIV, suggesting higher synchronization of activity in these networks compared to the healthy networks (Fig. 1D, table S1).

**Fig. 1.**
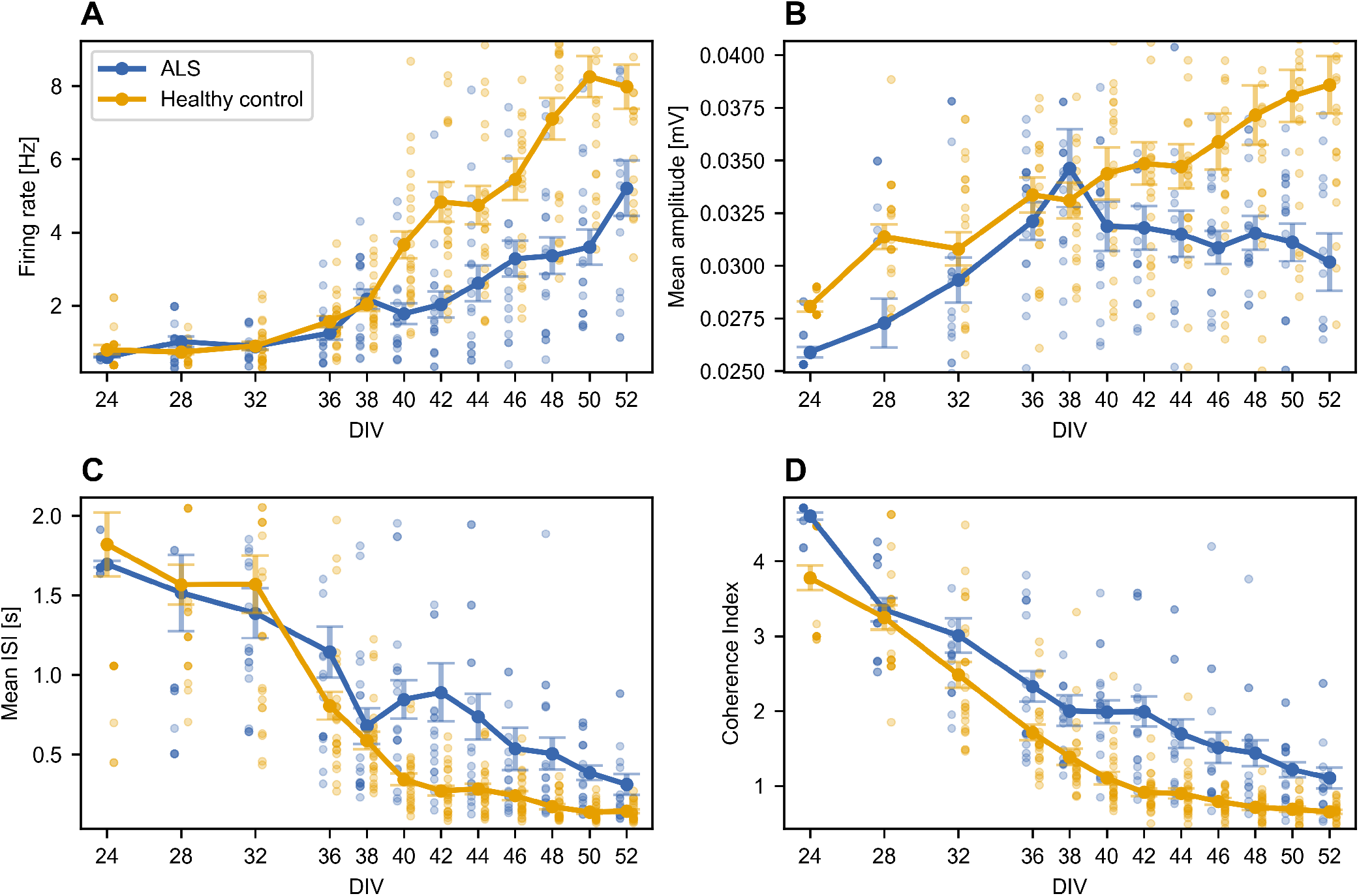
ALS motor neuron networks display functional deviations from controls upon network maturation. (**A**) Firing rate in Hz, (**B**) Mean amplitude of spikes in mV, (**C**) mean ISI in seconds, and (**D**) coherence index. The individual data points (mean from each MEA network in each group) are represented by the shaded circles, and the mean values for all networks in each group are represented by the solid lines and circles. Error bars show the SEM. Legend shared between figures.

### ALS networks exhibit reduced bursting

Both ALS and healthy networks showed signs of functional maturation over time, characterized by an increasing number of bursts, longer burst duration and shorter inter-burst intervals. The burst frequency was lower in the ALS networks compared to the controls, but this only became evident after 38 DIV (Fig. 2A). Significant differences emerged at 40 DIV (p<.001), and the burst frequency remained lower in the ALS networks until the final recording at 52 DIV, but no significant differences were identified at 46 DIV (p=.095917) and 52 DIV (p=.31792; table S1). There were large variations in the mean burst duration at 24 and 28 DIV, likely due to a low number of networks exhibiting bursting at these early time points (Fig. 2B). From 32-38 DIV, there were no significant differences between the ALS networks and the healthy networks, whereas from 40 DIV onward, the mean burst duration was significantly lower in the ALS networks. The mean IBI appeared to stabilize for both the ALS and the healthy networks from 36 DIV, but was slightly higher in the ALS networks (Fig. 2C). Significant differences were identified at 40-42 DIV and 46-50 DIV (table S1). The fraction of spikes in bursts was characterized by the same large variations as for the mean burst duration at 24 and 28 DIV (Fig. 2D). From 32-38 DIV, the fraction of spikes in bursts was slightly higher in the ALS networks compared to the healthy networks, albeit not significantly. From 40 DIV onward, the fraction of spikes in bursts was significantly lower in the ALS networks (table S1).

**Fig. 2.**
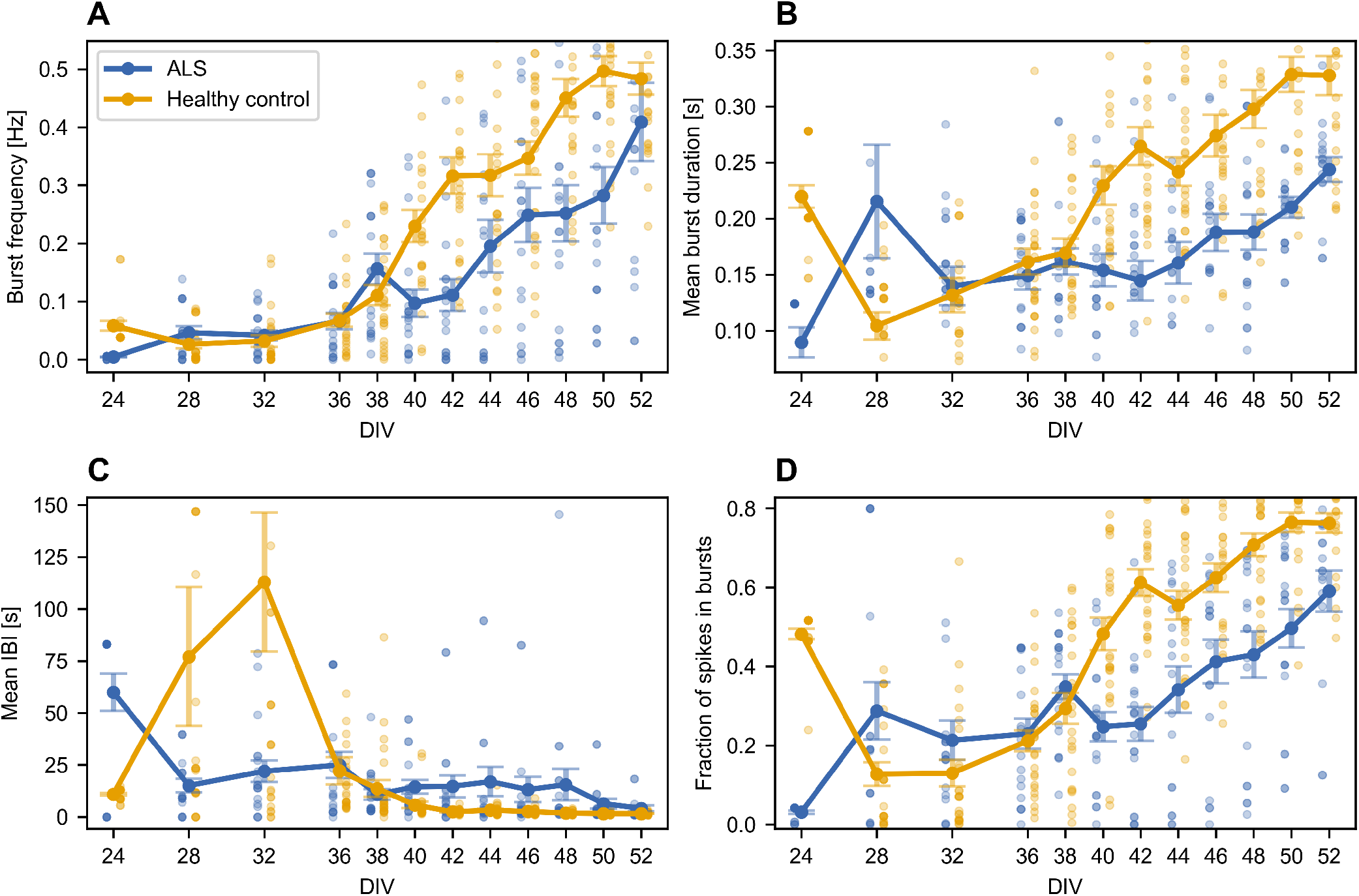
Characterization of bursting properties in ALS and healthy control neural networks. (**A**) Burst frequency in Hz, (**B**) Mean burst duration in seconds, (**C**) mean inter-burst-interval (IBI) in seconds, (**D**) fraction of spikes in bursts. The shaded circles show the individual data points, and the mean values for all networks in each group are represented by the solid lines and circles. Error bars show the SEM. Legend shared between figures.

### Impaired neurite outgrowth and branching in ALS motor neuron networks

To get a better understanding of the relationship between the structure of the neural networks and their functional development, immunocytochemistry assays were performed at three different time points: 18 DIV (one week after plating), 38 DIV (the day before overt functional differences between the groups started to emerge) and 52 DIV (the day of the final recording). In addition to confirming the presence of motor neuron-specific markers ISL1 (fig. S1), ChAT (fig. S2) and HB9 (Fig. 3), cytoskeletal marker expression also reflected different stages of maturation that corresponded to the functional development observed over time. Microtubule-associated protein 2 (MAP2) confirmed a more dispersed layer of neurons at 18 DIV (fig. S1A), with fewer dendritic trees, compared to 38 DIV (fig. S1B) and day 52 (fig. S1C). At 38 and 52 DIV, the motor neuron somata had clustered more together, and showed clearer dendritic branching. Expression of GFAP confirmed the presence of astrocytes, and showed a shift in the abundance of astrocytic processes from 18 DIV(fig. S2A) to 38 and 52 DIV (fig. S2B-C), suggesting that changes in astrocyte morphology in the presence of neurons are dependent on the duration in co-culture. The expression of neurofilament heavy (NFH) also confirmed the structural development of motor neuron networks from evenly dispersed neurons with short connections at 18 DIV (Fig. 3A) into highly clustered neuronal populations connected by longer and thicker axon bundles as they age (Fig. 3B-C).

**Fig. 3.**
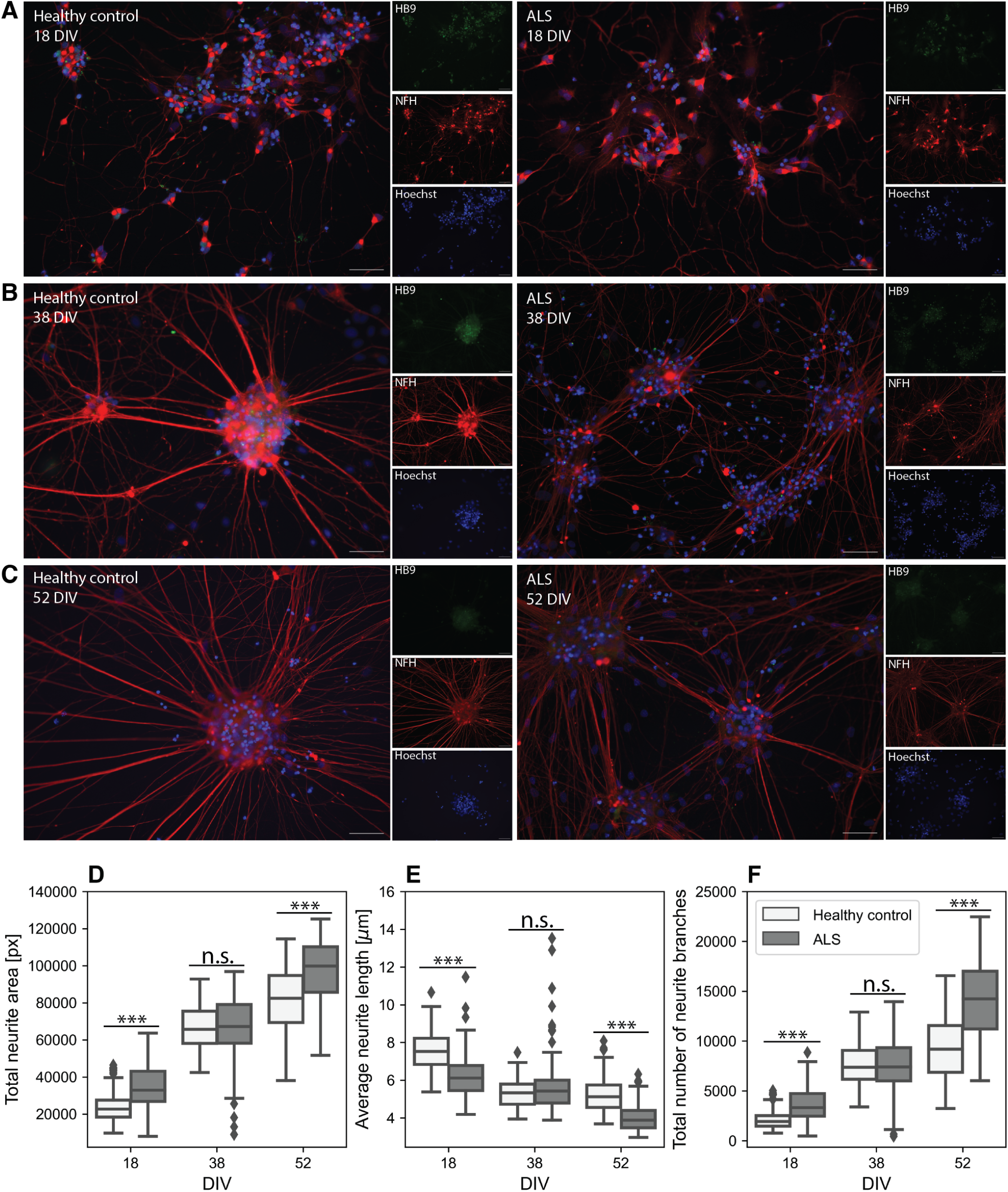
Structural development of motor neuron networks reveals increasing complexity. Immunocytochemistry assays confirmed positive expression of Neurofilament Heavy and HB9 at 18 DIV (**A**), 38 DIV (**B**) and 52 DIV (**C**). Scale bars: 100 *µ*m. (**D-F**) Box plots of quantification of neurite outgrowth showing the total area covered by neurites (D), the average neurite length (E), and the total number of neurite branches (F). Mann Whitney U-test. *** p<.001. n.s., not significant. Legend shared between D-F.

Although the ALS networks showed the same structural hallmarks of neuronal network maturation as the healthy networks, quantification of neurite length and density revealed subtle morphological changes in the ALS networks. The total area covered by neurites was significantly higher in the ALS networks at 18 (p<.001) and 52 DIV (p<.001) compared to controls, indicating higher neurite density in the ALS networks at these time points (Fig. 3D). The average neurite length however, was shorter in the ALS networks at 18 (p<.001) and 52 DIV (p<.001; Fig. 3E). The ALS networks also exhibited increased neurite branching compared to controls, again at 18 (p<.001) and 52 DIV (p<.001; Fig. 3F). These results suggest that ALS motor neurons exhibit enhanced neurite outgrowth and branching, resulting in a higher overall density in their networks, despite their neurite length being shorter. Interestingly, no significant differences were found between the groups at 38 DIV in the total neurite area, the average neurite length or the total number of branches. Additionally, the neurite length and branching in the healthy networks appeared to stabilize from 38 DIV to 52 DIV compared to the ALS networks. In fact, no significant difference from 38 DIV to 52 DIV was identified in the average neurite length in the healthy group (p=0.4885062; Fig. 3E), whereas the ALS group changed significantly (p<.001).

### RNA sequencing reveals aberrant expression of synaptic transcripts in ALS

We next sought to investigate the molecular underpinnings of the structural and functional differences observed in the ALS networks with transcriptomic analyses. Principal component analysis (PCA) of variance transformed gene counts showed clear segregation between the healthy group and the ALS group, but also a high degree of similarity between individual samples within each group (Fig. 4A). The different time points were clearly distinguishable, especially between 18 DIV and 38 DIV. Due to the clear segregation both between the ALS and the healthy group, and between individual time points, we hypothesized that their transcriptomic profile developed differently over time. We therefore first focused on the DEGs upon maturation, and performed GO enrichment analysis on ranked lists of DEGs from 18 DIV to 52 DIV in the healthy group (Fig. 4B) and the ALS group (Fig. 4C) separately. Many of the significantly enriched GO terms were conserved between the groups and were as expected involved in neuronal structures, development and signaling (Fig. 4D-E). Nevertheless, several ontologies identified in the healthy group were not present in the ALS group, including “axon”, “dendrite” and “dendritic tree”. On the other hand, “glutamatergic synapse” was found to be enriched in the ALS group and not in the healthy group, alongside extracellular matrix-associated ontologies.

**Fig. 4.**
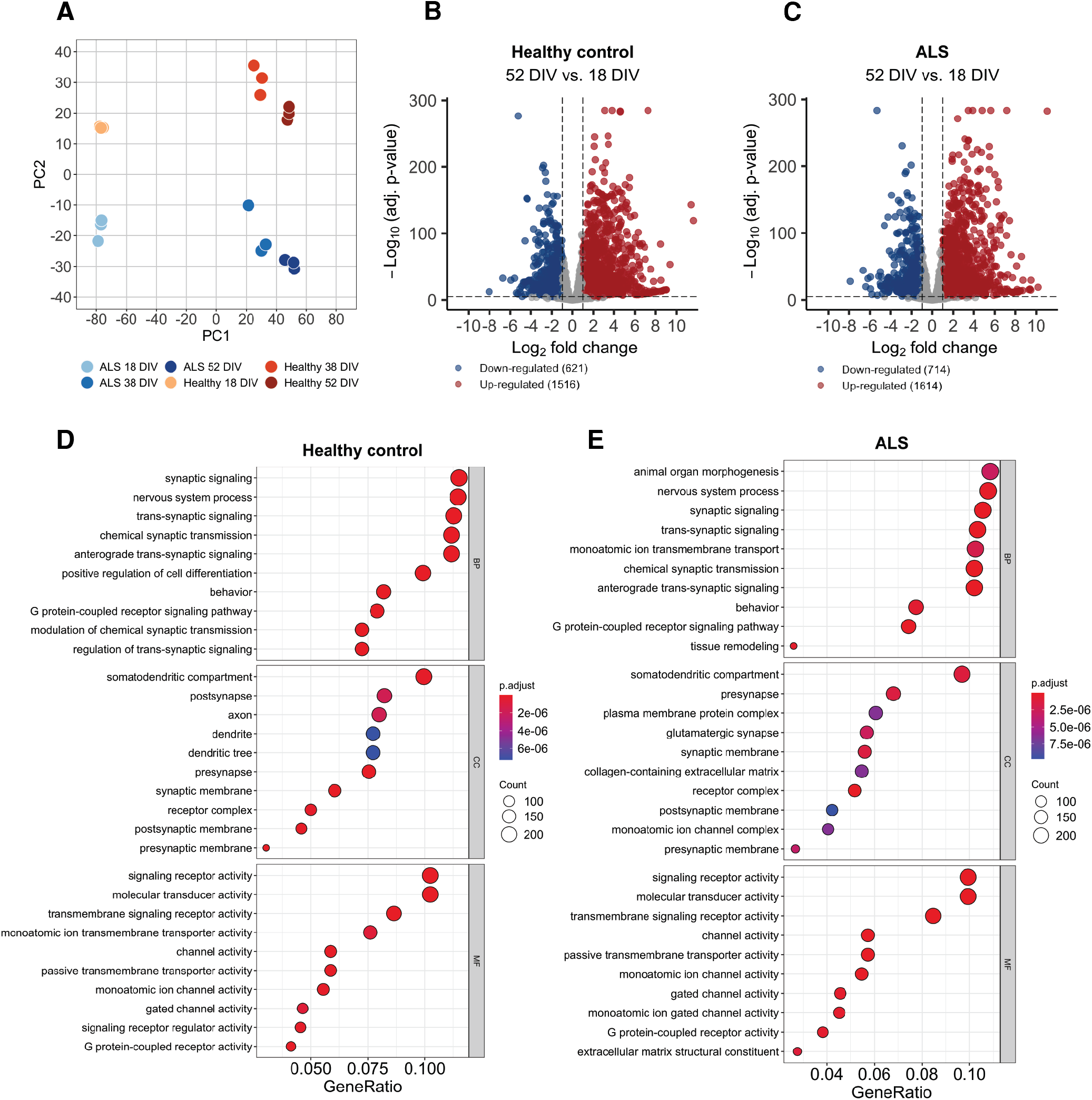
Transcriptional profiling of healthy and ALS motor neuron networks. (**A**) PCA of variance transformed gene counts in the ALS group and healthy group at each time point. Dots with the same color represent individual replicates within each group and time point. (**B**) Volcano plot of the upregulated (red) and downregulated (blue) DEGs from 18 DIV to 52 DIV in the healthy group. Dashed lines indicate thresholds for Log2 fold change (1.0; vertical lines) and the adjusted p-value (<.00001; horizontal lines). Legend shows the number of up- and downregulated genes. (**C**) Volcano plot of the upregulated (red) and downregulated (blue) DEGs from 18 DIV to 52 DIV in the ALS group. (**D**) Dot plot of the top ten enriched GO terms for “Biological Process” (BP), “Cellular Component” (CC) and “Molecular Function” (MF) in the healthy group. The color of the dots illustrates the adjusted p-value with warmer color reflecting more significant values, and dot size represents the gene count (i.e. number of genes) associated with a specific term. (**E**) Dot plot of the top ten enriched GO terms for “Biological Process” (BP), “Cellular Component” (CC) and “Molecular Function” (MF) in the ALS group.

We then compared the transcriptomes of the ALS and the healthy group at each individual time point, i.e. 18 DIV, 38 DIV and 52 DIV. Volcano plots revealed differential expression of several cell-adhesion proteins including *PCDH5, PCDHGA3, PCDHGA7* and *PCDHA13* at all three time points (Fig. 5A-C). To get a better understanding of how gene expression differences developed over time, we identified the top 30 DEGs at each individual time point (Fig. 5D-F). This confirmed the involvement of protocadherins and revealed some stable expression patterns over time, including dysregulation of *KHDRBS2, MYO1G, PTPRT* and *NLRP2*. Nevertheless, some alterations in the top 30 DEGs were observed from timepoint to timepoint, including increased downregulation of the cell-adhesion and migration gene *AJAP1* from 18 DIV (Fig. 5D) to 38 DIV (Fig. 5E). These findings thus suggested dysregulation of several cell-adhesion proteins in the ALS group, including protocadherins, which play important roles in axonal, dendrite and synapse development and maintenance (28).

**Fig. 5.**
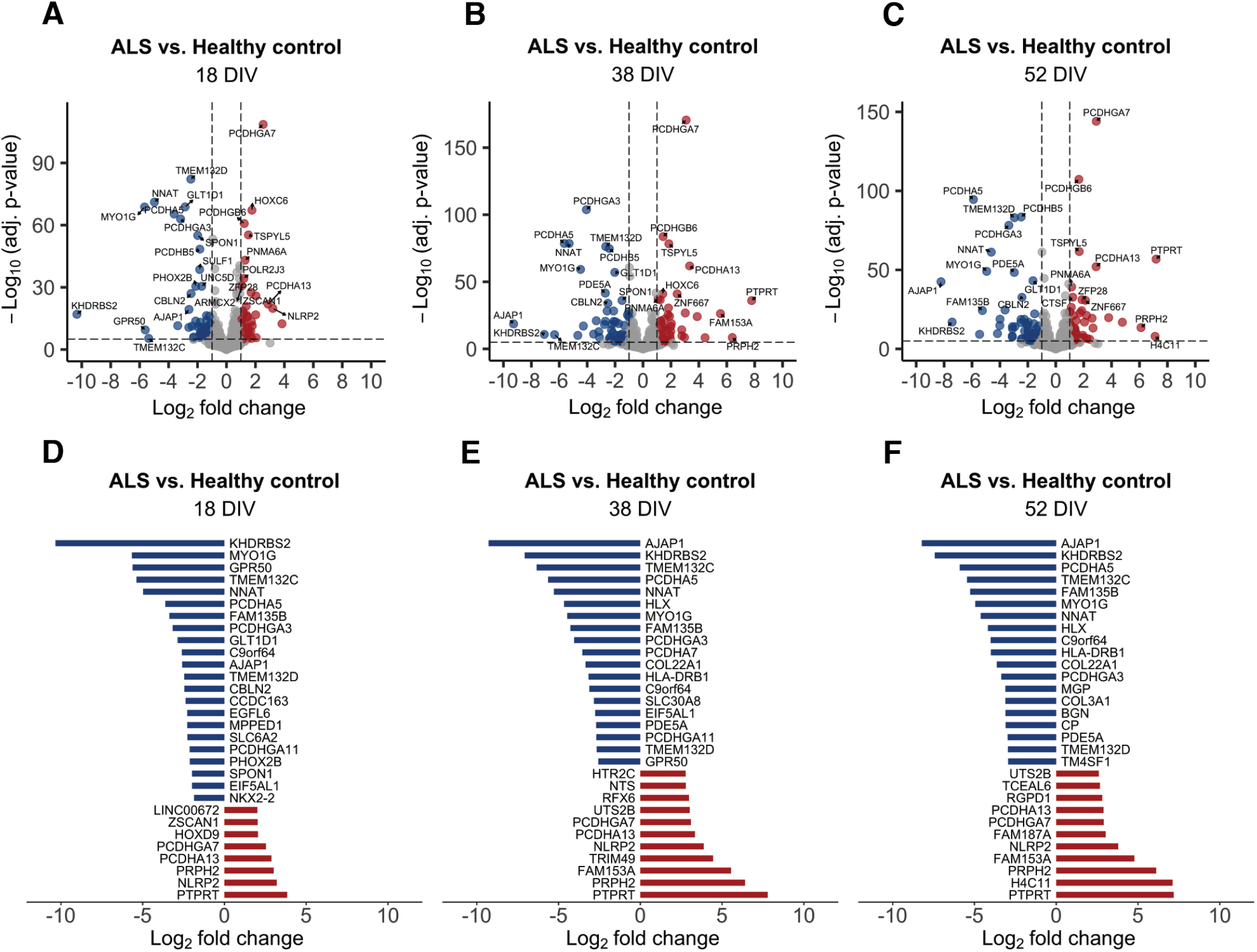
Differential gene expression between healthy and ALS motor neuron networks. (**A-C**) Volcano plots of upregulated (red) and downregulated (blue) DEGs in the ALS group compared to the healthy group at 18 DIV (A), 38 DIV (B) and 52 DIV (C). Genes with highest Log2 fold change and/or adjusted p-value are named. Dashed lines show thresholds for Log2 fold change (1.0; vertical lines) and the adjusted p-value (<.00001; horizontal lines). (**D-F**) Bar plots showing the top 30 DEGs in the ALS group compared to the healthy group at 18 DIV (D), 38 DIV (E) and 52 DIV (F). X-axis shows the Log2 fold change. Negative values represent downregulated genes, and positive values represent upregulated genes.

Considering the observed differences in neuronal activity and the importance of stable and functional synapses for efficient neuronal signaling, we focused on the synaptic genes in further analyses. GO enrichment analysis of the synaptic gene lists demonstrated that although several ontologies were shared between the healthy (Fig. 6A) and the ALS group (Fig. 6B), most of the Cellular Component ontologies enriched in the healthy group were absent in the ALS group. Further differences included enrichment of “neurotransmitter receptor activity” in the healthy group only, and enrichment of catecholamine transport and secretion in the ALS group only. To investigate the specific pathways and genes that could underlie these differences, volcano plots of differentially expressed synaptic genes between the ALS and the healthy group were established for 18 DIV (Fig. 6C), 38 DIV (Fig. 6D) and 52 DIV (Fig. 6E). Several genes involved in synaptic transmission, neurite outgrowth and synaptic plasticity were found to be dysregulated in the ALS group. For example, *CBLN2*, which is involved in maintenance of excitatory synapses, was consistently downregulated in the ALS group at all three time points. *FRMPD4*, also required for maintenance of excitatory synaptic transmission, was upregulated at 18 and 38 DIV. *SHISA6*, involved in glutamatergic excitatory synaptic transmission, was upregulated at both 18 DIV and 52 DIV. NMDA-receptor subunit 3A, *GRIN3A*, was downregulated at 52 DIV. Aberrantly expressed genes involved in neurite outgrowth included *RGS14*, which was downregulated at all time points, and *NTNG2* and *SLITRK4*, which were upregulated at 18 DIV and 38 DIV, respectively. Additionally, the synaptic plasticity gene *PRSS12* was upregulated at 38 DIV, and both *PRSS12* and *ARC* were upregulated at 52 DIV. Finally, *PTPRT*, involved in signal transduction and cell adhesion, was upregulated at all three time points, and the log2 fold change and significance level increased over time. In summary, these transcriptomic profiling data revealed dysregulation of multiple pathways in ALS motor neuron networks, including synaptic transmission and maintenance, neurite outgrowth and cell adhesion.

**Fig. 6.**
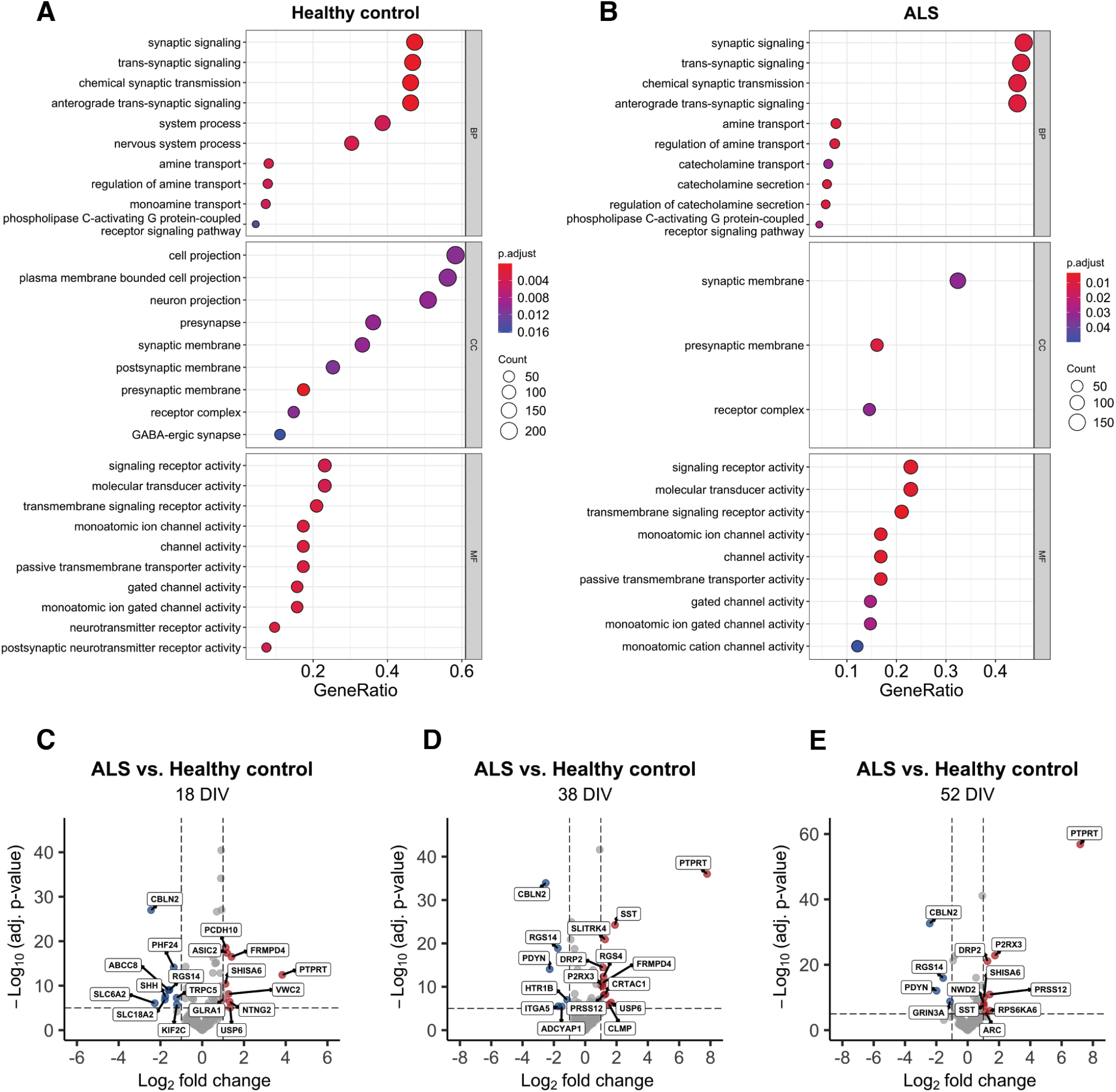
Transcriptional analysis unravels dysregulated pathways associated with synapses in ALS motor neuron networks. (**A-B**) Dot plots of the top ten enriched GO terms in the synaptic genes for “Biological Process” (BP), “Cellular Component” (CC) and “Molecular Function” (MF) in the healthy (A) and ALS group (B). (**C-E**) Volcano plots demonstrating upregulated (red) and downregulated (blue) synaptic genes in the ALS group at 18 DIV (C), 38 DIV (D) and 52 DIV (E).

## Discussion

In this study, we conducted longitudinal systematic recordings of hiPSC-derived motor neurons with a confirmed endogenous ALS mutation. We identified specific time points for the emergence of network dysfunction, specifically reduced firing rate and amplitude, impaired bursting, and higher synchrony of network activity. These changes were accompanied by structural alterations. We show that ALS motor neurons have increased neurite density and branching but shorter neurite length, indicating aberrant neurite outgrowth. These dysfunctions appear to result from synaptic alterations, as revealed by transcriptional analyses, suggesting an intrinsic failure in the ALS networks to establish and maintain functional synapses over time. Importantly, although the differences in the transcriptome were present throughout the experiment, the functional consequences of the associated differences appeared to be time dependent. This was demonstrated by the clear differences observed in neural network firing rate and bursting only after 38 DIV (Fig. 1 and Fig. 2), and by the temporary stabilization of structural impairments at 38 DIV, as shown in Fig. 3. Crucially, the ALS networks remained viable and functional over time, suggesting that impaired synaptic function does not directly cause motor neuron degeneration, but may instead initiate network-wide compensatory mechanisms.

Several recent studies have identified synaptic impairment in ALS patient-derived neurons, suggesting that synaptic dysfunction may be an early event in the pathophysiological cascade (20, 29, 30, 25, 31). One study identified disruptions in genes involved in synaptic transmission and synaptogenesis including *CBLN1, CBLN2, CBLN4* and *SLITRK2* in C9-ALS patient-derived motor neurons (20). Similarly, Perkins et al. (29) identified altered expression of *CBLN1* and genes involved in cell-cell adhesion, including the *γ*-protocadherin *PCDHGC4* in C9-ALS patient-derived cortical neurons. Both cerebellins and protocadherins are crucial for synaptic stabilization and maintenance (32, 28), indicating an important role for these families of genes in maintaining synaptic integrity in ALS. In striking accordance with these previous findings, we also observed altered gene expression of *CBLN2, SLITRK4* (Fig. 6) and several protocadherins (Fig. 5) in the ALS motor neurons. Notably, these patterns were stable throughout our study, suggesting that synaptic instability occurs prior to functional deficits. This is also supported by previous work reporting disruptions in synaptic stability and transmission before cell death (33). We did not directly assess neuronal death in the present study, but previous studies have not observed substantial cell death in hiPSC-derived motor neurons with ALS pathology (25, 31). Furthermore, indirect measures of cell death such as a marked reduction in firing rate or detachment of neurons did not reveal any clear signs of cell death. It is therefore unlikely that the structural and functional differences observed in this study were caused by neuronal loss. Instead, our study corroborates previous evidence indicating that synaptic maintenance and stability are affected early in ALS (33, 25, 31) and may over time result in structural and functional alterations.

The altered expression of genes involved in synaptic maintenance, neurite outgrowth and cell adhesion in the ALS networks may have resulted in aberrant neurite branching, as indicated by the increased neurite density and branching in the ALS networks (Fig. 3D-F). Axon pathfinding and target selection require accurate integration of attractant and repellent guidance cues, thus the dysregulation of such cues, including protocadherins (28), *SLITRK4* (34) and *NTNG2* (35) may have resulted in inaccurate target specificity in the ALS networks. Aberrant branching may reflect an underlying deficit in establishing and maintaining synapses. Importantly, we have previously observed aberrant neuritic outgrowth in a hiPSC-derived Parkinson’s disease model (36). Aberrant branching has also been identified in mutant TDP-43 overexpressing zebrafish models (37, 38) and human ALS patient-derived motor neurons (39, 40), underscoring the possible involvement of aberrant branching in neurodegenerative diseases. Interestingly, we also observed reduced average neurite length in the ALS networks (Fig. 3E), in correspondence with previous investigations in human C9orf72 mutant motor neurons (30). This may have affected the ALS networks’ capacity to efficiently establish long-range connections, resulting in compensatory aberrant branching and suboptimal structural network connectivity.

Axon pathfinding and neurite branching play crucial roles in the development of the central nervous system, but increased neurite branching may indicate underlying pathology in later stages when the network is established. Notably, we observed differences in neurite density, branching, and average neurite length between the ALS and the healthy networks at the first (18 DIV) and the last time point (52 DIV), demonstrating that these features were not only present during the networks’ initial development. Instead, they may reflect intrinsic impairment affecting the structural network. Interestingly, no difference was observed between the two groups at 38 DIV in the total neurite area, the average length or the number of branches. This was also accompanied by a temporary upregulation of SLITRK4 at 38 DIV, which is involved in inhibition of neurite outgrowth (41), suggesting initiation of compensatory mechanisms. These findings are further supported by the observation that the average neurite length stabilized in the healthy networks from 38 to 52 DIV, but not in the ALS networks. Additionally, no differences were observed in network activity at 38 DIV, followed by a clear reduction in firing rate and bursting compared to healthy networks at later time points (Fig. 1), indicating the presence of temporary compensation to preserve function.

Aberrant neurite branching and synapse instability could affect signal propagation efficiency within the network, resulting in a decreased firing rate and less bursting, as observed in the electrophysiology data from 40 DIV onward (Fig. 1 and Fig. 2). We also found a lower spike amplitude in the ALS networks which appeared to stabilize from 40-52 DIV compared to healthy networks. Lower spike amplitude in ALS patient-derived motor neuron networks is consistent with previous reports (25, 26). Importantly, reduced spike amplitude results in lower Ca^2+^ influx (42), which has been shown to reduce bursting (43), consistent with the bursting profile identified in the ALS networks.

Bursting is an essential property of efficient information processing in in vitro neural networks, and closely related to synchronous neural activity (18). Interestingly, we observed higher synchrony of network activity in the ALS networks, as indicated by the coherence index (Fig. 1D), despite the reduced bursting frequency at the electrode level. Thus, while the ALS networks exhibited less bursting at the local level compared to their healthy counterparts, the overall network activity was more synchronized, indicating initiation of compensatory mechanisms to sustain the connections and information integration across the network. Increased network burst frequency has previously been reported in ALS patient-derived cortical neurons, combined with shorter network burst duration (29). Excessive synchrony may however also result in impaired local segregation, i.e. less functional specialization (44). This may render the network less flexible and adaptable, ultimately resulting in reduced resilience to perturbations. In fact, it has been shown that functional brain networks in ALS patients reorganize into a more connected and synchronized network as disease progresses, thereby maintaining functionality while also becoming increasingly vulnerable to additional pathological processes (8). In contrast, reduced synchrony of network activity was also observed in ALS patient-derived motor neuron networks on high-density MEAs, despite an increase in network bursting and vulnerability to perturbations (26). As such, the exact relationship between bursting dynamics and synchrony, and their role in early stages of ALS pathogenesis require further investigation.

A denser network structure is often associated with higher neuronal firing rate (18). Indeed, a considerable amount of work has shown increased firing and hyperexcitability in ALS patients (6, 11), animal models of the disease (45, 46) and hiPSC-derived models (21), including our own previous work (26). Nevertheless, a growing body of evidence indicates that hyperexcitability is a transient phenotype followed by hypoexcitability (23, 24, 25, 47). The hypoexcitable phenotype has been attributed to an imbalance in Na^+^ and K^+^ currents (23, 22). In a computational motor neuron degeneration model, it has been shown that reduced availability of ATP may lead to reduced activity of the Na^+^/K^+^ ATPase pumps and loss of Na^+^ and K^+^ ion gradients, resulting in chronic membrane depolarization (48). Experimental evidence supporting this has been provided in both an hiPSC-derived model (23) and a mouse model (15), both showing a more depolarized resting membrane potential compared to controls. This has been suggested to contribute to reduced excitability by increasing the sodium inactivation and potassium activation (15).

The structural and functional deficits observed in the present study can be explained within an energy homeostasis perspective: impaired synapse stability may result in an increased need for ATP to maintain the existing synapses, and to account for increased branching in an effort to establish more connections. Insufficient ATP production due to mitochondria and axonal transport dysfunction (30) may in turn result in loss of ion gradients and altered resting membrane potential. Computational modeling studies have demonstrated that altered membrane potential in response to reduced energy availability can result in a dynamic shift from a transient hyperactive or hyper-synchronized state to a hypoexcitable state (48, 49). Importantly, we did not observe early hyperexcitability in our study, in contrast with some previous reports (23, 24, 25, 47), possibly due to methodological differences such as electrophysiological methods, the presence of astrocytes, and the frequency of media changes. Still, it is notable that the reduced firing rate only emerged after 38 DIV, a finding that is in accordance with other studies reporting hypoexcitability in hiPSC-derived neurons first after four weeks (25, 24). Furthermore, downregulation of neuronal activity in hyperactive neurons (21) and upregulation of activity in hypoactive neurons (22, 33) have been reported to have neuroprotective effects, underscoring the importance of the intrinsic electrophysiological state of the neurons for treatment outcomes. In summary, our findings demonstrate the importance of spatiotemporal dynamics on pathological processes, and this may also have implications for the timing of therapeutic interventions.

The aberrant neurite branching and altered network dynamics observed in the ALS patient-specific neural networks in this study can be considered compensatory responses to intrinsic synaptic dysfunction. It would be interesting to further investigate the role of synaptic impairments in ALS with strategies aimed at restoring synaptic function and network balance. Such interventions should also be expanded to cases with other ALS-associated genetic mutations, as well as sporadic cases. It must also be taken into consideration that the exact function and interplay between many genes are not fully understood (50). As such, the implications of dysregulated gene expression patterns for neural network function also require further investigation, especially in relation to therapeutic modulation of gene expression. Importantly, we observed variations over time in all analyses, thereby emphasizing the dynamical nature of neural network behavior and the impact investigation time points and duration can have on results. Such aspects may be of particular relevance in hiPSC-derived disease models, as their functional development still lacks characterization (27, 51), especially in pathological states. Furthermore, network dynamics may not only be influenced by the underlying genotype but also epigenetic factors such as donor age, a feature which may be better captured with direct reprogramming techniques bypassing the stem cell stage (52). These dynamics should also be considered when evaluating the efficacy and ideal time windows for therapeutic approaches in the future.

In conclusion, this study provides evidence for early synaptic dysfunction in ALS, accompanied by dynamic structural and functional impairments and compensation. Such mechanisms may place the motor neuron networks in a metabolically demanding state, thereby rendering the networks vulnerable to other ongoing pathological processes ultimately converging in neurodegeneration. Stabilizing synaptic integrity is therefore a highly relevant approach for restoring neural network function and slowing disease progression in ALS.

## Materials and Methods

### Cell lines and motor neuron differentiation

Human iPSCs derived from fibroblasts from a confirmed ALS patient (female, 64, C9orf72 mutation) and a healthy control donor (female, 49) were expanded on vitronectin-coated (A14700, Thermo Fisher) 100 mm petri dishes using complete mTeSR Plus medium (mTeSR Plus Basal medium and mTeSR Plus 5X supplement; 100-0276, StemCell Technologies). The medium was exchanged daily until the hiPSCs reached approximately 80% confluence, at which the cells were differentiated into motor neurons according to the section “Prodecure E” in the protocol published by Nijssen et al. (53). Briefly, the iPSCs were dissociated into a single-cell suspension using TrypLE Express (12604021, Thermo Fisher), and resuspended in N2/B27 medium, consisting of the indicated mixture described in (53). The cells were kept in the incubator (37°C, 5% CO2) on an orbital shaker throughout the embryoid body (EB) stage, i.e. the first 10 days of the protocol. The medium was freshly supplemented every day according to table S2. On day 10 of the motor neuron differentiation protocol, the EBs were dissociated with TrypLE Express, and resuspended in B27 medium supplemented with the day 10 factors.

### Neural networks

On differentiation day 10, motor neurons were seeded on Axion Biosystems 24-well CytoView MEA plates (M384-tMEA-24W). The C9orf72 motor neurons and the healthy control motor neurons were seeded on one plate each, resulting in n = 24 neural networks for each group. The plates were pre-coated with Poly(ethyleneimine) solution (PEI; P3141, Merck) and natural mouse Laminin (23017015, Thermo Fisher). 70 *µ*L of 0.05% PEI diluted in HEPES (15630-056, Thermo Fisher) were added to each well, carefully concentrated around the electrode area, and incubated over night. The next day, the PEI was removed and the wells were rinsed four times with distilled water. All the water was removed and the plates were left to air dry over night. On the following day, Laminin was diluted in PBS (D8537, Sigma-Aldrich) to a final concentration of 20 *µ*g/mL, and the wells were coated with 70 *µ*L for at least one hour prior to seeding. When seeding, the Laminin was removed and a small volume (30 *µ*L) containing 30,000 motor neurons were plated within the electrode area in each well (1310 neurons/mm^2^). The plates were left in the incubator for 1 hour to allow the cells to attach before adding more medium to a final volume of 500 *µ*L per well. The media was fully replaced on day 11, 12 and 13 of the differentiation protocol (53).

On day 13, human astrocytes at P4 (K1884, Thermo Fisher Scientific Gibco) were seeded in each well. The astrocytes were thawed and centrifuged according to the manufacturer’s instructions, and resuspended in day 13 motor neuron medium. All media on the MEA plates were removed and replaced with 400 *µ*L fresh day 13 medium, before dropwise adding the remaining 100 *µ*L containing 3000 astrocytes to each well (10% of the motor neuron population). From day 13 onward, the neural networks were maintained with full media changes every other day for the remainder of the experiment. The timeline for establishment of neural networks on MEAs is shown in Fig. 7.

**Fig. 7.**
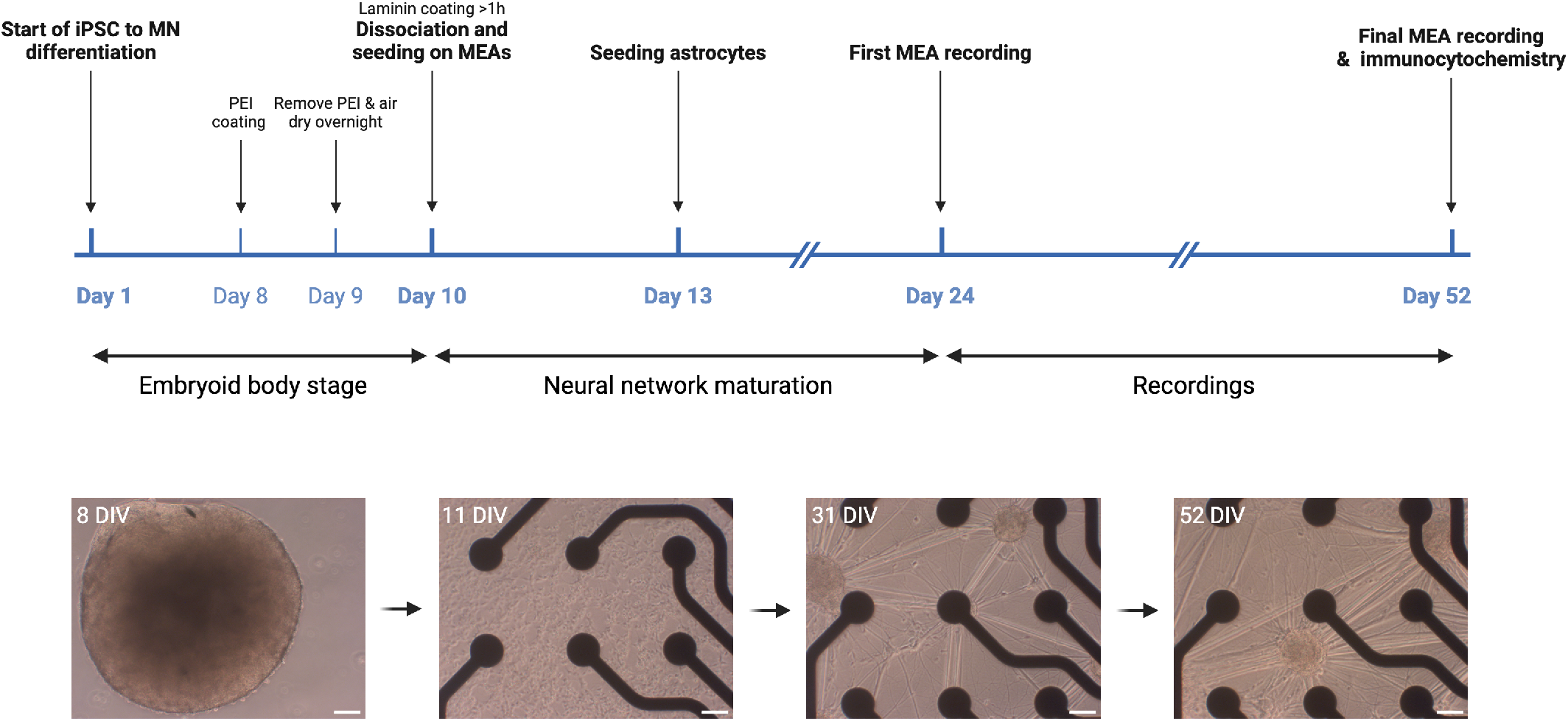
Experimental timeline for electrophysiology experiments. Motor neuron differentiation was initiated when iPSCs reached approximately 80% confluency. During the first ten days of the differentiation, the cells were kept on an orbital shaker to facilitate embryoid body formation. After ten days, the embryoid bodies were dissociated and plated onto precoated MEAs. Three days later, human astrocytes were plated on the MEAs, and at 24 DIV the first recording was performed. The final MEA recording was performed at 52 DIV, and cells for immunocytochemistry assays were fixed on the same day. Phase-contrast images show the development of an example network from the embryoid body stage at 8 DIV, to a motor neuron network on an MEA at 11 DIV, 31 DIV and 52 DIV. Scale bars: 50 *µ*m. Figure created with BioRender.com.

### Electrophysiology

Recordings of neural activity were made using the Axion Maestro Pro MEA system (Axion BioSystems, GA, USA), which has a built-in incubator ensuring stable temperature and CO_2_ conditions (37°C, 5% CO2). Data acquisition was done with the AxIS Navigator software (version 3.6.2) with a sampling frequency of 12.5 kHz. The plates were left for 30 minutes before starting a recording to allow activity to stabilize after being moved to the recording platform. Each recording lasted for 20 minutes and the first recording was made at 24 DIV. To monitor the activity, subsequent recordings were made every four days until 36 DIV, at which the networks started to display more stable activity. Between 36-52 DIV, recordings were made every other day, resulting in a total of 12 recordings per group. In the recording at 28 DIV for the ALS networks, the last 19.5 seconds of the 20 minute recording were not recorded due to a fault in the Maestro Pro system. Spikes were detected by the AxIS Spike Detector, using an adaptive threshold crossing of 7 standard deviations of the continuous data stream within frequency band 200Hz-3kHz.

### Network data analysis

Data was processed using the Python programming language (Python Software Foundation) with the Pandas package. Statistical analysis was performed using the packages Scipy (54) and Pingouin (55), and figures were generated using Seaborn and Matplotlib.

To ensure that the analyzed networks had coherent activity throughout the entire recording, electrodes with a firing rate lower than 0.2 Hz and wells with a well firing rate below 0.5 Hz were excluded from network analysis. These numbers were chosen as they filtered out the networks with very low firing rate, without excluding all of the data points at the earliest recording time points. The total number of networks included in the analysis at each time point is reported in table S3 in the Supplementary file.

Analysis of electrophysiology data was based on (56). The firing rate *f* was defined as *f* = (*spikes −* 1)*/n*Δ*t*, where *spikes* is the total number of recorded spikes in a given well, *n* is the number of electrodes considered to be active throughout the recording and Δ*t* is the time difference between the first and the last recorded spike.

The instantaneous firing rate *f*_*window*_ was calculated for parameters using shorter time windows, i.e. the coherence index, and was defined as *f*_*window*_ = *spikes/n*Δ*t*. Here, spikes is the total number of spikes in the given time window, *n* is the number of active electrodes and Δ*t* is the width of the given window. The instantaneous firing rate was calculated as a moving window with window size of 100 ms and step size of 10 ms, resulting in partially overlapping windows.

The inter-spike interval (ISI) was defined as the time interval between two consecutive spikes. Bursts were defined as a train of at least four spikes with an ISI less than 100 ms. The burst frequency *f*_*burst*_ was defined as *f*_*burst*_ = *bursts/n*Δ*t*, with bursts being the total number of bursts throughout the recording in a given well and Δt the time difference between the first and the last recorded spike. The inter-burst-interval (IBI) was defined as the time interval between two consecutive bursts. Based on these definitions, the mean duration of bursts and the fraction of spikes in bursts were identified.

Due to a low number of active electrodes in some of the networks, network burst detection was not performed. Instead, the coherence index was computed as an alternative approach to assess synchronization of neural network activity. The coherence index was defined as the ratio between the standard deviation and the mean of the instantaneous firing rate (57). A higher coherence index indicates a more synchronous network activity, whereas lower synchrony is reflected by a firing rate with little oscillation.

### Bulk mRNA sequencing

Time points for mRNA sequencing analyses were identified based on observed differences between the ALS and the healthy group in the MEA data. The time points chosen were 18 DIV, 38 DIV and 52 DIV. This allowed us to investigate potential differences between the groups before, during and after the emergence of overt differences in neural network behavior. Co-cultures of motor neurons and astrocytes were plated and maintained on six-well plates according to the same procedures described above. Samples were collected from three independent cultures from each cell line at the three time points. RNA was extracted using the Qiagen RNeasy Mini Kit (74104; Qiagen, Hilden, Germany), according to the manufacturer’s instructions. Quality control, library preparation and sequencing was performed at the Genomics Core Facility at NTNU. RNA quality was assessed using Agilent 2100 Bioanalyzer (Agilent Technologies Inc., Santa Clara, CA, United States) and quantity determined using a NanoDrop spectrophotometer. Sequencing libraries were generated with Illumina Stranded mRNA Prep kit and sequencing of 2×50 bp paired-end reads was performed on an Illumina NovaSeq 6000 (Illumina Inc., San Diego, CA, United States). The sequences were aligned to the human GRCh38 reference genome using STAR (58). Salmon was used for quasi-mapping to quantify transcript abundance (59). The R package tximport (60) was used to generate the associated gene counts and the gene counts were normalized. Differentially expressed genes (DEGs) were identified using DESeq2 (61). The analysis pipeline is available at the GitHub rna-seq repository (62).

### Transcriptome data analysis

Data analysis and visualization was performed in RStudio (RStudio IDE 2023.12.0+369; (63)). Gene Ontology (GO) enrichment analysis was performed using clusterProfiler (version 4.10.0; (64)) with the org.Hs.eg.db annotation package (version 3.18.0) on lists of DEGs with thresholds Log_2_ fold change≥1 and Benjamini-Hochberg adjusted p-value<.05, and plotted using enrichplot (65). Synaptic genes were extracted from the DEG lists using the SynGO geneset tool (66). Volcano plots were generated using EnhancedVolcano (67) to visualize highly significant DEGs, with thresholds Log_2_ fold change≥1 and Benjamini-Hochberg adjusted p-value<.00001. All other plots were generated using ggplot2 (68).

### Immunocytochemistry and imaging

For immunocytochemistry assays, motor neurons and astrocytes were co-cultured on 8-well chamber slides in parallel with MEAs and 6-well plates for mRNA sequencing. Cells were fixed at day 18, day 38 and day 52 with 3% glyoxal solution, based on the protocol by (69). The solution consisted of 71% MQ water, 20% ethanol absolute, 8% glyoxal (128465, Sigma-Aldrich) and 1% acetic acid (1.00063, Sigma-Aldrich). After 15 minutes, the glyoxal was removed and the samples were washed three times with PBS (D8662, Sigma-Aldrich) for five minutes. The final PBS wash was removed and 0.5% Triton-X (1.08643, Sigma-Aldrich) diluted in PBS was added to each well for five minutes to permeabilize the cells. The samples were subsequently washed two times with PBS for 5 minutes, before adding a blocking solution consisting of 5% Goat Serum (G9023, Sigma-Aldrich) diluted in PBS. The chips were left in room temperature for 1 hour on an orbital shaker at 30 rpm. The blocking solution was then replaced with PBS containing 5% goat serum and the primary antibodies (table S4) and the chips were stored on a tilting shaker in a cold room (3 °C) until the next day.

The following day, the samples were washed three times with PBS for five minutes each, before adding the secondary antibodies (table S4). The chips were kept on an orbital shaker at 30 rpm in room temperature for three hours. Nuclei were stained with Hoechst diluted 1:2000 in PBS (bisbenzimide H 33342 trihydrochloride; 14533, Sigma-Aldrich) for 10 minutes, before washing three times with PBS for five minutes. The samples were then washed once with distilled water. Two types of chamber slides were used, of which one type had removable wells (177402, Thermo Fisher). These were mounted on glass cover slides using anti-fade fluorescence mounting medium (ab104135, abcam). The chamber slides with non-removable cover slips (80807, ibidi GmbH) were filled with fresh water and stored in a cold room. Imaging was done using an EVOS M5000 microscope (Invitrogen) with an Olympus UPLSAPO, 20x/0.75 NA (N1480500) objective, and the following LED light cubes: DAPI (AMEP4650), GFP (AMEP4651), TX-Red (AMEP4655) and CY5 (AMEP4656). Image processing was performed in Fiji v1.54f.

### Image quantification of neurite outgrowth

Quantification of neurite length and density was done using the NeuroConnectivity macro set (70, 71) in Fiji. Neural networks immunolabeled with Neurofilament Heavy and Hoechst were used for analysis and imaged using an EVOS M7000 microscope (Invitrogen) with an EVOS LWD 20x/0.45 NA (AMEP4982) objective, and LED light cubes DAPI (AMEP4650) and TX-Red (AMEP4655). Images were acquired using an automated scan protocol capturing 15% of the well area in a random order, yielding a total of 56 images per well. The autofocus setting was used to achieve optimal focus at each capture. Pixel sizes were 0.309 *µ*m x 0.309 *µ*m. Four wells were imaged per group at three time points, i.e. 18 DIV, 38 DIV, and 52 DIV. After acquisition, images were manually assessed and images with artifacts (out of focus, edge of a well or debris) were excluded from further analysis as this could interfere with the neurite segmentation. The total number of images included in the analysis per group were as follows: Healthy 18 DIV, n = 185; Healthy 38 DIV, n = 166; Healthy 52 DIV, n = 184; ALS 18 DIV, n = 170; ALS 38 DIV, n = 151; ALS 52 DIV, n = 198. Neurite detection was based on (72) and nuclei were detected using the Stardist algorithm (73). The Python packages Scipy (54), Pingouin (55) and Scikit-posthocs (74) were used for statistical comparisons.

### Statistical analysis

The Shapiro-Wilk’s test was applied to test for normality in each group at each individual time point. Statistical comparisons between the groups were assessed using a two-sample independent Welch’s t-test when the data was normally distributed, and a Mann Whitney U-test in the case of non-normality. Comparisons within each group across time points in the neurite outgrowth quantifications were assessed using the Kruskal-Wallis test, followed by Conover’s test for multiple comparisons with Bonferroni correction. Results were determined as significant if p<.05.

## Acknowledgments

Bulk mRNA sequencing was provided by the Genomics Core Facility (GCF), Norwegian University of Science and Technology (NTNU). GCF is funded by the Faculty of Medicine and Health Sciences at NTNU and Central Norway Regional Health Authority.

## Author contributions

Conceptualization: AMK, AS, IS

Methodology: AMK, NC, AS, IS

Investigation: AMK

Formal Analysis: AMK, NC

Visualization: AMK, NC

Funding acquisition: AS, IS

Project administration: AS, IS

Supervision: AS, IS

Writing – original draft: AMK

Writing – review & editing: AMK, NC, AS, IS

## Supplementary Materials

The Supplementary Materials file contains:

Fig. S1 to S2

Table S1 to S4

