## Supplementary Materials for "Dysregulation of synaptic transcripts underlies network abnormalities in ALS patient-derived motor neurons"

Kollstrøm et al.

Ioanna Sandvig:

This file includes:

Fig. S1 to S2
Tables S1 to S4

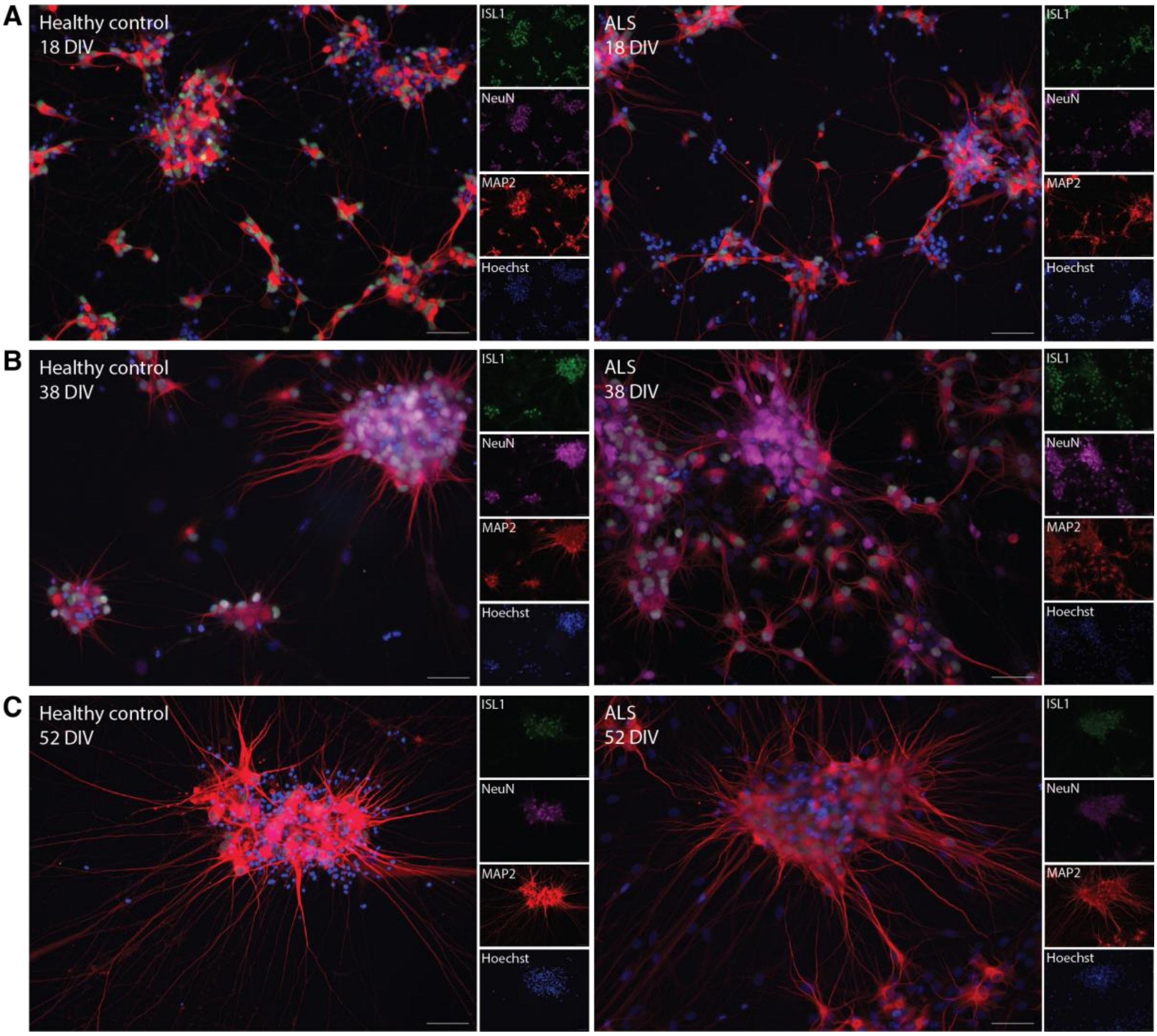

*Fig. S1. Motor neuron networks derived from hiPSCs expressed the motor neuron-specific marker Islet-1, and neuronal markers MAP2 and NeuN. Example images from a healthy (left) and ALS network (right) at 18 DIV (**A**), 38 DIV (**B**) and 52 DIV (**C**). Scale bars: 100 µm.*

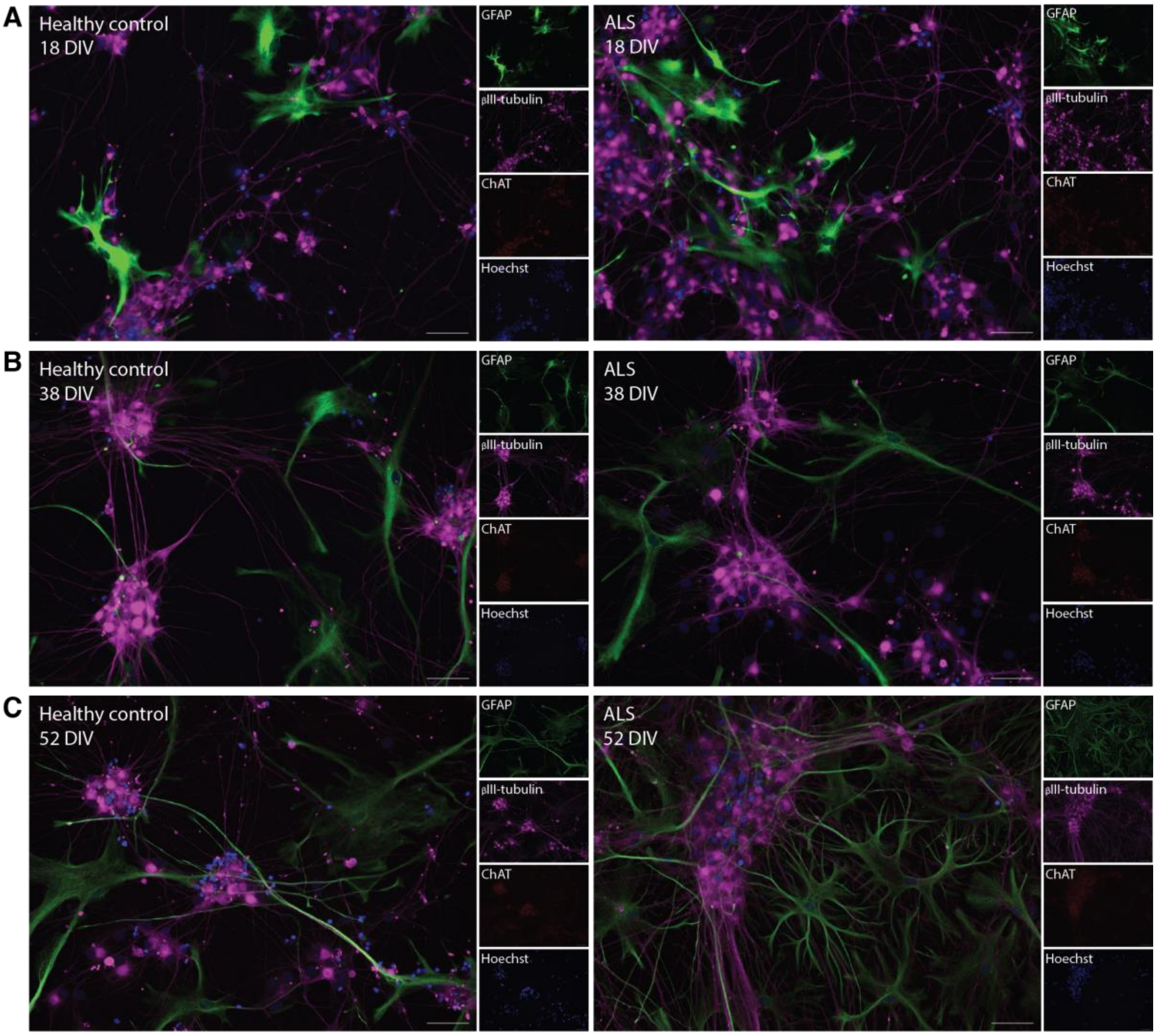

*Fig. S2. Immunocytochemistry confirmed the presence of astrocytes in the neural networks with positive expression of GFAP, as well as expression of βIII tubulin and the motor neuron-specific marker ChAT. Scale bars: 100 µm.*

*Table S1. P-values from statistical comparisons of ALS vs. healthy control for the network parameters used in the MEA analysis. Welch's t-test or Mann-Whitney U-test were applied as appropriate. Significant values (p <.05) in bold.*

| | **Day** | 24 | 28 | 32 | 36 | 38 | 40 | 42 | 44 | 46 | 48 | 50 | 52 |
|---|---|---|---|---|---|---|---|---|---|---|---|---|---|
| **Network parameter** | | | | | | | | | | | | | |
| Firing rate [Hz] | | 0.80437 | 0.294137 | 0.54885 | 0.097001 | 0.651857 | **0.00014** | **0.000053** | **0.005196** | **0.006083** | **0.000112** | **0.000005** | **0.007619** |
| Mean amplitude [mV] | | **0.000015** | **0.022888** | 0.166823 | 0.537623 | 0.598889 | 0.111536 | **0.042618** | **0.041427** | **0.004972** | **0.001469** | **0.000108** | **0.000204** |
| Mean ISI [s] | | 0.80437 | 0.294137 | 0.54885 | 0.097001 | 0.835266 | **0.00014** | **0.000053** | **0.005196** | **0.013197** | **0.000112** | **0.000005** | **0.014544** |
| Coherence index | | **0.000006** | 0.838066 | **0.041039** | **0.011969** | **0.000916** | **0.000023** | **0.000002** | **0.000095** | **0.000106** | **0.000018** | **0.000046** | **0.000388** |
| Burst frequency [Hz] | | **3.951622e-08** | 0.204806 | 0.138733 | 0.635384 | 0.276289 | **0.000424** | **0.000035** | **0.039788** | 0.095917 | **0.001703** | **0.000544** | 0.31792 |
| Mean burst duration [s] | | **3.852850e-08** | 0.068948 | 0.207292 | 0.794255 | 0.398645 | **0.00245** | **0.000016** | **0.001033** | **0.001256** | **0.000027** | **1.020522e-07** | **0.000252** |
| Mean IBI [s] | | **0.014883** | **0.038932** | 0.079985 | 0.924448 | 0.797269 | **0.00309** | **0.001934** | 0.123835 | **0.045996** | **0.00024** | **0.030837** | 0.219197 |
| Fraction of spikes in bursts | | **3.951622e-08** | 0.185997 | 0.114471 | 0.924448 | 0.282837 | **0.000121** | **8.803486e-08** | **0.008735** | **0.004972** | **0.000288** | **0.000012** | **0.00208** |

*Table S2. Overview of cell media components used for motor neuron differentiation. Adapted from Nijssen et al. (2019).*

| Catalog number & supplier | Reagents | Days 1-2 | Days 3-9 | Day 10 | Days 11-12 | Day 13 onward |
|---|---|---|---|---|---|---|
| 31331028, Thermo Fisher | DMEM/F12 + GlutaMAX | 50% | 50% | | | |
| 21103049, Thermo Fisher | Neurobasal | 50% | 50% | 100% | 100% | 100% |
| 17502048, Thermo Fisher | N2 (100X) | 0.5X | 0.5X | | | |
| 17504044, Thermo Fisher | B27 (50X) | 0.5X | 0.5X | 1X | 1X | 1X |
| 15070063, Thermo Fisher | Penicillin/Streptomycin | 1X | 1X | 1X | 1X | 1X |
| Y0503, Sigma-Aldrich | Y-27632 | 5 µm | | 5 µm | | |
| 1614, Tocris | SB-431542 | 40 µm | | | | |
| 6053, Tocris | LDN-193189 | 200 nM | | | | |
| SML1046, Sigma-Aldrich | CHIR-99021 | 3 µm | | | | |
| R2625, Sigma-Aldrich | All-*trans* retinoic acid | | 200 nM | 200 nM | | |
| 4366, Tocris | SAG | 500 nM | 500 nM | | | |
| A4403, Sigma-Aldrich | L-Ascorbic acid | 200 µm | 200 µm | 200 µm | 200 µm | 200 µm |
| 2634, Tocris | DAPT | | | 10 µm | 10 µm | |
| 450-10, Peprotech | GDNF | | | 10 ng/mL | 10 ng/mL | 10 ng/mL |
| 450-02, Peprotech | BDNF | | | 10 ng/mL | 10 ng/mL | 10 ng/mL |

*Table S3. Overview of the number of neural networks included in network analysis for each group from 24 to 52 DIV.*

| DIV | 24 | 28 | 32 | 36 | 38 | 40 | 42 | 44 | 46 | 48 | 50 | 52 |
|---|---|---|---|---|---|---|---|---|---|---|---|---|
| ALS | 3 | 8 | 14 | 14 | 14 | 16 | 16 | 15 | 15 | 15 | 14 | 14 |
| Healthy control | 4 | 12 | 19 | 22 | 23 | 24 | 24 | 24 | 24 | 24 | 24 | 24 |

.

*Table S4. Overview of antibodies and concentrations used for immunocytochemistry.*

| Primary antibodies | Concentration | Catalog number and supplier |
|---|---|---|
| Rabbit anti-Islet 1 | 1:250 | EP4182, abcam |
| Rabbit anti-HB9 | 1:200 | ab221884, abcam |
| Rabbit anti-GFAP | 1:5000 | ab7260, abcam |
| Mouse anti-NeuN | 1:1000 | ab279295, abcam |
| Mouse anti-Beta III Tubulin | 1:1000 | ab14545, abcam |
| Chicken anti-Neurofilament heavy | 1:1000 | ab4680, abcam |
| Chicken anti-MAP2 | 1:5000 | ab5392, abcam |
| Chicken anti-ChAT | 1:200 | ab34419, abcam |
| Secondary antibodies | Concentration | Catalog number and supplier |
| Goat anti-rabbit Alexa Fluor 488 | 1:1000 | A11008, Fisher Scientific |
| Goat anti-mouse Alexa Fluor 647 | 1:1000 | A21236, Fisher Scientific |
| Goat anti-chicken Alexa Fluor 568 | 1:1000 | ab175477, abcam |
